# Cellular stress modulates severity of the acute respiratory distress syndrome in COVID-19

**DOI:** 10.1101/2022.09.09.507257

**Authors:** Gustavo Rico-Llanos, Óscar Porras-Perales, Sandra Escalante, Daniel Vázquez, Lucía Valiente, María I. Castillo, José Miguel Pérez-Tejeiro, David Baglietto-Vargas, José Becerra, José María Reguera, Ivan Duran, Fabiana Csukasi

**Affiliations:** Networking Biomedical Research Center in Bioengineering, Biomaterials, and Nanomedicine (CIBER-BBN), Andalusian Centre for Nanomedicine and Biotechnology, Málaga, Spain.; Laboratory of Precision Medicine in Musculoskeletal and Inflammatory Diseases, IBIMA-Bionand Platform, Malaga, Spain.; Infectious Disease Unit, Hospital Regional de Malaga, Malaga, Spain.; Department of Cell Biology, Genetics, and Physiology, Faculty of Sciences, University of Málaga, Málaga, Spain.; Veterinary clinic of exotic pets ARACAVIA, Málaga, Spain.; Networking Biomedical Research Center in Neurodegenerative Disease (CIBER-NED), Madrid, Spain.; Institute for Memory Impairments and Neurological Disorders, University of California, Irvine, California.; Department of Orthopaedic Surgery, University of California-Los Angeles, Los Angeles, CA 90095, USA.

**Keywords:** COVID-19, *acute respiratory distress syndrome* (ARDS), binding-immunoglobulinprotein (BiP/GRP78/HSPA5), cytokine storm, cell surface BiP/GRP78, cellular stress, TNF, 4-PBA.

## Abstract

Inflammation is a central pathogenic feature of the acute respiratory distress syndrome (ARDS) in COVID-19. Previous pathologies such as diabetes, autoimmune or cardiovascular diseases become risk factors for the severe hyperinflammatory syndrome. A common feature among these risk factors is the subclinical presence of cellular stress, a finding that has gained attention after the discovery that BiP (GRP78), a master regulator of stress, participates in the SARS-CoV-2 recognition. Here, we show that BiP serum levels are higher in COVID-19 patients who present certain risk factors. Moreover, early during the infection, BiP levels predict severe pneumonia, supporting the use of BiP as a prognosis biomarker. Using a mouse model of pulmonary inflammation, we demonstrate that cell surface BiP (cs-BiP) responds by increasing its levels in leukocytes. Neutrophiles show the highest levels of cs-BiP and respond by increasing their population, whereas alveolar macrophages increase their levels of cs-BiP. The modulation of cellular stress with the use of a clinically approved drug, 4-PBA, resulted in the amelioration of the lung hyperinflammatory response, supporting the anti-stress therapy as a valid therapeutic strategy for patients developing ARDS. Finally, we identified stress-modulated proteins that shed light into the mechanism underlying the cellular stress-inflammation network in lungs.

## Introduction

The COVID-19 pandemic has challenged our understanding of the inflammatory response. COVID-19 is an infectious disease that becomes severe and lethal through a poorly known mechanism whose output is barely prognosed by risk factors and comorbidities^1, 2^. Since the first wave in 2020, we have learned that the *severe acute respiratory syndrome coronavirus 2* (SARS-CoV-2) is able to induce a hyperinflammatory response commonly known as *cytokine storm*, with consequences that are very similar to other diseases with a *cytokine release syndrome* (CRS). Although COVID-19 is considered a systemic disease, the respiratory system is the most affected, where the CRS is better defined as *acute respiratory distress syndrome* (ARDS).

A major problem with COVID-19 has been the inability to predict which patients could develop a severe disease and the most accurate method to predict the outcome of the infection has been the measurement of interleukin-6 (IL-6)^1, 3–5^. However, IL-6 can only be detected after the development of acute symptoms, leaving clinical risk factors as our only way of prognosis ^6, 7^. Risk factors correlated with COVID-19 include age (median > 62), gender (with increased tendency in men), and chronic pathologies such as diabetes, chronic liver disease, hypertension, immunodeficiency, chronic obstructive pulmonary disease (COPD), smoking history, among others^8–11^, and while they have been useful for early follow-up of symptoms, they are not accurate predicting severity and they do not include subclinical manifestations that also cause severe CRS.

The cytokine profile of COVID-19 has been studied since the beginning of the pandemic concluding that it does not differ much from other forms of ARDS and sepsis^1^. There is plenty of evidence that elevated levels of different cytokines like IFN-γ, IL-6, IL-1β, IL-10 and MCP-1 are higher in COVID-19 patients. There is also a clear association between others such as IP-10, MCP-1, MIP-1α, TNF-α and IL-6 and COVID-19 severity when comparing ICU-patients with non-ICU patients^1, 3–5^. However, they do not predict the severity outcome of the disease and they cannot be used reliably to explain why some patients develop a severe response to the infection.

The binding-immunoglobulin protein (BiP), also called Grp78, and encoded by the gene *Hspa5*, is an endoplasmic reticulum (ER) chaperone that acts as a master regulator of the unfolded protein response (UPR) and ER-stress signaling pathways^12, 13^. Increased levels of BiP have been found in several pathological conditions such as liver disease^14^, metabolic disorders and atherosclerosis^15, 16^, cardiovascular diseases^17^, diabetes^18^, cancer^19, 20^, acute lung injury (ALI)^21^, autoimmune disorders^22, 23^, different forms of subclinical inflammation^24, 25^, aging^26^ and neurodegenerative diseases^27^. Many of these pathologies are risk factors for COVID-19. Although the main fraction of BiP in the cell is dedicated to regulate the UPR and the secretory pathway, BiP has also been found to translocate to other compartments upon stress stimulus (cytoplasm, mitochondria, extracellular matrix and cell surface)^28, 29^. Cellular surface BiP (csBIP or csGRP78) acts as a co-receptor for different signaling pathways (PI3K, CD109, Cripto, CD44v, alpha2M, caspases 7 and 8 and clathrin dependent pathways) modulating cell proliferation, differentiation, survival and apoptosis^30, 31^. It is therefore considered a key protein in the crosstalk between multiple signaling pathways, working as a sensor of various cellular stresses to maintain homeostasis^20^. Moreover, BiP has been found to participate in many viral infections including COVID-19^32, 33^, Ebola, Zika, Dengue, Japanese Encephalitis Virus, Coxsackievirus A9, Borna Disease Virus and the Middle-East Respiratory Syndrome coronavirus (MERS)^33–39^. However, even when there is solid evidence that dysregulated levels of both intracellular and csBiP are linked to these diseases, much work is needed to fully understand the mechanism by which this protein modulates inflammation in response to the stress signals that increase its expression or promote its localization to the cell membrane. Nonetheless, BiP is a multifunctional chaperone that goes beyond the ER compartment when the cell is under any type of cellular stress (infection, hypoxia, heat shock, ER and oxidative stress)^40–44^.

After BiP was found to act as a co-receptor of Angiotensin converting enzyme-2 (ACE2) for SARS-CoV-2 virus^33^ BiP has been hypothesized to favor virus entry into the cell, however, evidence from other pathologies in which BiP acts as co-receptor indicate that this role goes beyond virus recognition or replication. For example, BiP has been related as an immunomodulatory factor interacting with the Jak/STAT system and possibly with other cytokine intracellular signaling components, including IL-6^45, 46^. From all this evidence, our group and others have suggested that BiP and the cellular stress must have a modulatory effect on the hyper-inflammatory response produced after infection with SARS-CoV-2 and its clinical outcome^47, 48^.

Here, we investigated the role of cellular stress and BiP in the modulation of the ARDS inflammatory response in samples from COVID-19 patients and a mouse model of ARDS. We demonstrate that BiP levels correlate with the severity of ARDS. Furthermore, we show that the localization of BiP on the cell surface is increased in the immune cell lineages during ARDS proportionally to the severity of the inflammatory response and identify a network of proteins that mediate this pathological process. Our results support the use of BiP as a prognosis biomarker of severe pneumonia and offer a new therapeutic strategy for diseases with ARDS such as COVID-19.

## Results

### BiP levels in blood serum correlate with COVID-19 comorbidities and severity

Besides being a SARS-CoV-2 coreceptor, BiP is increased in several pathologies identified as risk factors of COVID-19, however, no study has investigated the connection of BiP with the risk factors of severe COVID-19. To correlate BiP with COVID-19 severity we measured BiP levels in blood serum from 194 patients of the first wave of the pandemic (March-June 2020), obtained at the beginning of the SARS-CoV-2 infection during the first medical evaluation. All patients were confirmed PCR-positive. This cohort included patients with different degree of clinical severity, from asymptomatic to lethal COVID-19. Thirty healthy blood donors, without infection or any detectable pathology, were used as a control of BiP levels (see supplementary document 1 about blood donor selection/exclusion criteria). We established that 95% of the healthy control population has levels of BiP in serum below 181 pg/ml. Thus, we considered high levels of BiP those above 181 pg/ml. The average BiP level was higher in patients compared to control although it did not reach significance (*P* value = 0.0789). We detected high levels of BiP in the blood serum of 35 out of the 194 COVID-19 patients (18.04%) (Figure 1A). To determine which risk factors and comorbidities were present in patients with increased BiP, we analyzed how BiP levels correlated with 43 clinical parameters (Figure 1 and S1). BiP levels were higher in male patients and individuals above 60 years old, a group particularly vulnerable to suffer severe COVID-19 disease (Figure 1B-C). Among previous conditions, BiP was also elevated in patients with a history of hypertension, diabetes, immunosuppression and previous respiratory pathologies (Figure 1D-G). Within previous respiratory pathologies we were able to determine that increased BiP levels in blood correlated specifically with previous history of chronic obstructive pulmonary disease (COPD) (Figure 1H). These results indicate that increased BiP levels correlate to several risk factors of COVID-19 patients with a strong significance with the presence of previous respiratory pathologies.

**Figure 1.**
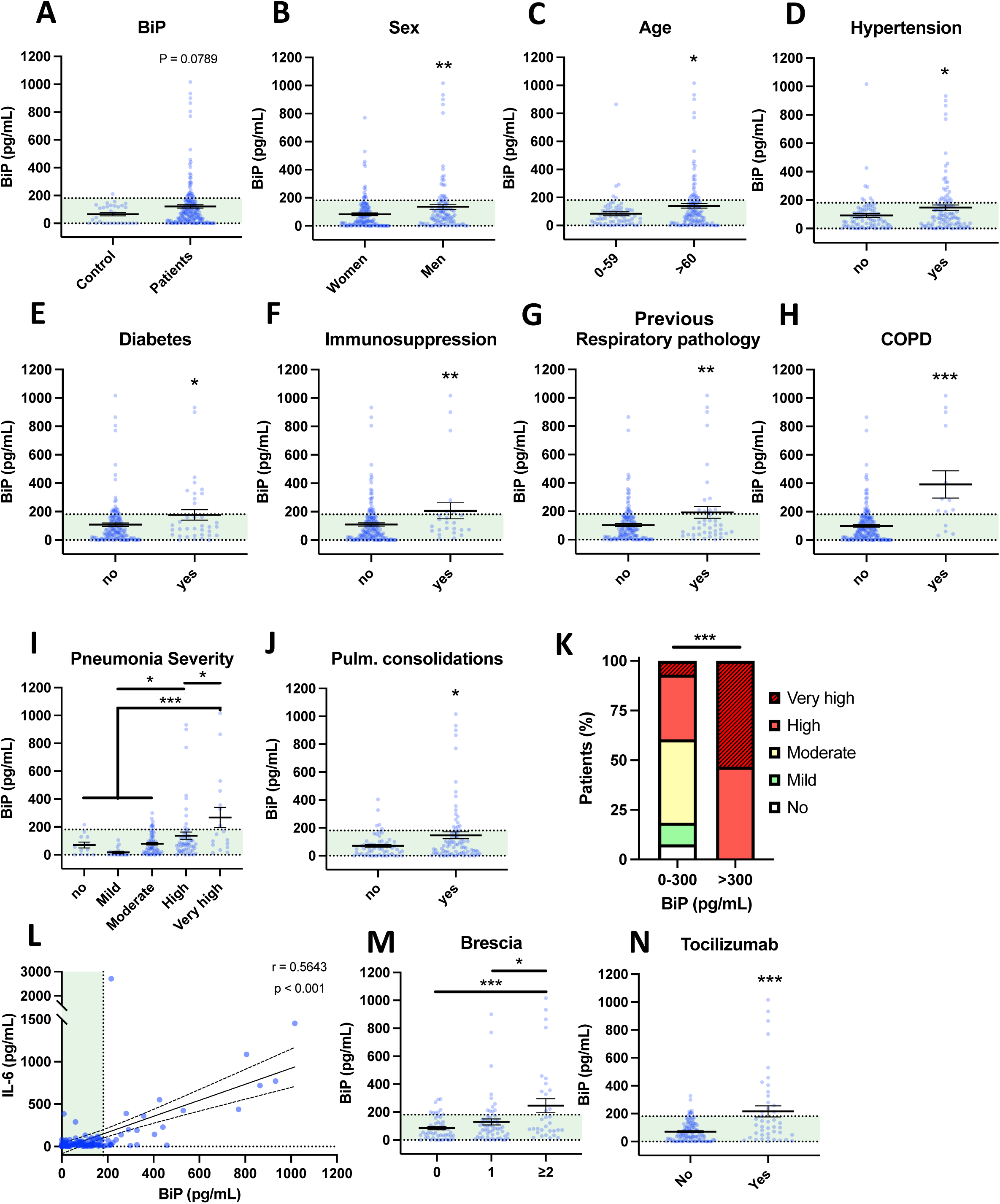
Serum BiP levels are increased in certain groups of COVID-19 patients. **(A-L)** Serum BiP levels classified by group of patients/donors. Black lines and whiskers denote the mean ± SEM of every data set. Green areas represent normal BiP levels in serum (0 and 181 pg/mL, respectively) defined between 5^th^ and 95^th^ percentiles of healthy donor’s data set. **General** BiP levels in total cohort: healthy control patients (n=30) versus COVID-19 patients (n=194) **(A)**; BiP levels classified by Sex **(B)**, Age **(C)**, and previous comorbidities **(D-H)**. **(I)** Serum BiP in patients classified by pneumonia severity in 5 levels depending on oxygen saturation, Tachypnea and need for mechanic ventilation. **(J)** BiP levels analyzed by radiological presence of pneumonia pulmonary consolidations developed during COVID-19. **(K)** Stacked bar plot showing percentage of patients with BiP levels below or above the selected critical threshold (300 pg/mL) who developed severe pneumonia (denoted by color code in legend). **(L)** Scatter plot showing a positive correlation between BiP levels versus IL-6 levels in blood serum tested by Pearson’s correlation coefficient. **(M)** BiP levels analyzed by Brescia-COVID Respiratory Severity Scale. **(N)** BiP levels analyzed by application of Tozilizumab treatment. \**P* < 0.05, \*\**P* < 0.01, \*\*\**P* < 0.001 indicate statistical significant differences between indicated samples for a Two-Tailed unpaired t-Test **(A-H, J, N)**, One-Way ANOVA with a Tukey’s multiple comparisons test **(I,M)** and Chi-square test **(K)**.

To determine the predictive potential of BiP levels in blood we analyzed the relationship between BiP serum levels and respiratory parameters corresponding with a severe COVID-19 like development of pneumonia. To categorize severity in pneumonia, patients were clinically classified into 5 groups according with the need for oxygen saturation, tachypnea and mechanic ventilation (Table S1). We observed a solid correlation between BiP and pneumonia severity groups “high” and “very high” (Figure 1I), which includes patients with oxygen saturation below 90%, possible tachypnea and in need for mechanical ventilation. Supporting this correlation, BiP levels were also significantly elevated in patients presenting pulmonary consolidations, a radiological finding typical of severe pneumonia (Figure 1J). From this data, we determined the distinctive threshold of BiP levels above which all patients developed severe pneumonia under these two categories. Thus, any patient with BiP levels 300pg/ml or higher during the initial stages of the infection developed severe pneumonia and needed high flow mechanical ventilation (Figure 1K). These results suggest that serum levels of BiP are a useful biomarker of the severe pneumonia output.

Next, we studied the correlation between BiP and IL-6 levels, the most widely used inflammation and severity marker for COVID-19. Our data showed a significant correlation between systemic BiP and IL-6 (Figure 1L), which confirms not only the association between BiP and COVID-19 severity but also suggests a connection between cellular stress and inflammation in the COVID-19 mechanism of disease. No other relevant changes were observed in blood values in correlation with BiP serum levels (Figure S2).

To further evaluate the predictive character of BiP in serum, we compared BiP levels with severity indexes for COVID-19. Systemic BiP was correlated with COVID-19 severity measured by its specific scale: Brescia-COVID-19 Respiratory Severity Scale^49^ that scored respiratory fatigue, respiratory rate >22, PaO2 <65 mmHg, SpO2 <90% and significantly worsening Chest X-Ray. More precisely, BiP levels were significantly elevated in patients with a Brescia index ≥2 (Figure 1M). Interestingly, above this score, patients in our cohort were considered for Tocilizumab (Anti IL-6) treatment which accordingly correlated the selection criteria of high IL-6 with high levels of BiP in serum (Figure 1N). Given the association between BiP levels and respiratory parameters, we also analyzed other pneumonia scores such as Pneumonia Outcomes Research Team (PORT) or the Pneumonia Severity Score CURB65. However, while PORT showed a weak association to BiP levels, CURB65 showed no change regarding to the stress marker (Figure S2).

Altogether, these results indicate that the levels of BiP in serum, measured at the time of hospital admission, correlate with a variety of general pre-existing comorbidities and could be used as a biomarker of the severity output that is especially relevant in relation with respiratory pathologies.

### Treatment with 4-PBA ameliorates the severity of the hyperinflammatory response in ARDS

Given the association between the stress marker BiP and the cytokine IL-6, we next studied the connection between this UPR regulator and other markers of the inflammatory response to determine which factors could be modulated by cellular stress. As the respiratory conditions are among the most relevant correlations with the levels of BiP in serum, we used an inflammation mouse model of acute respiratory distress syndrome (ARDS) that consists on the intranasal administration of lipopolysaccharide (LPS) from *E. Coli*. To determine whether cellular stress is involved in the inflammatory response, we also studied the effect of the application of the molecular chaperone 4-PBA after LPS challenge, an approved drug for several pathologies^50–54^ that reduces cellular stress and inflammation^55–57^. Hemograms performed after administration of LPS revealed a systemic neutrophilia, lymphopenia and monocytosis, mimicking the human response to SARS-CoV-2 infection. 4-PBA treatment seemed to partially rescue the blood parameters, however, these changes where not statistically significant at the systemic level except for the monocyte numbers that increased with LPS and were significantly rescued with 4-PBA (Figure S3).

To study in depth the inflammatory response in lungs we measured 14 cytokines in the broncoalveolar lavage fluid (BALF), selected by its relevance in lung tissues during the COVID-19 and/or cytokine storm syndrome. Among these, changes in IL-6, IL-1β and TNF-α levels were the best documented in the hyperinflammatory response associated to COVID-19 and ARDS. LPS challenge induced a significant increase in all the cytokines included in this study (IL-1β, TNF-α, IL-6, IFN-γ, IL-17a, MIP-1α, MCP-3, GM-CSF, IP-10, RANTES, MIG, IL-18 and MCP-3) except for IL-12p70 whose increase was not statistically significant (Figure 2A-N). Nor PBS instillation (C-) neither 4-PBA alone induced any changes in the cytokine levels. These data validated our mouse model induced by LPS instillation and established a well-characterized response of acute lung inflammation (ALI) at the cytokine level.

**Figure 2.**
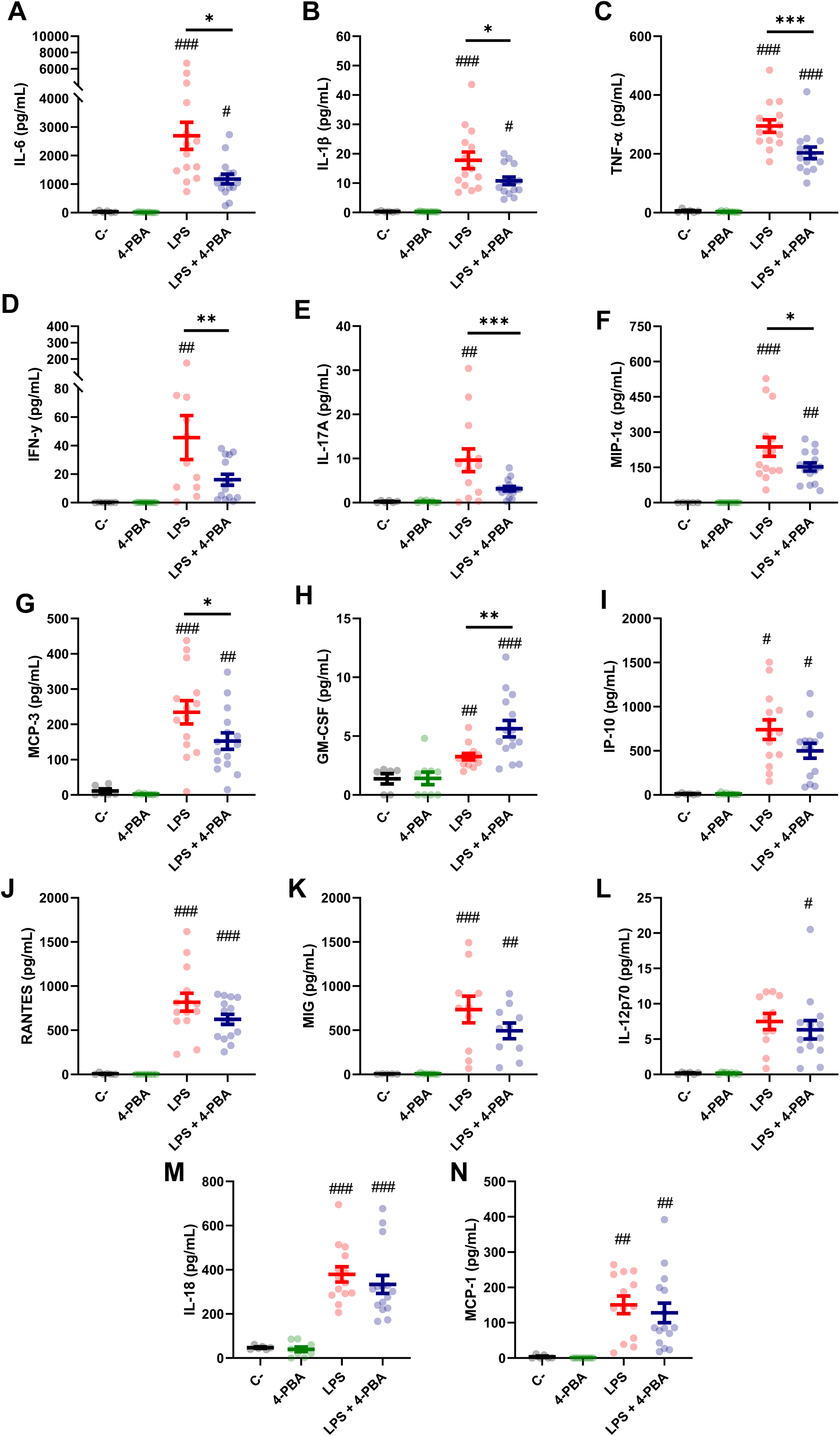
Bronchoalveolar cytokine profile after LPS challenge and 4-PBA treatment in ARDS model. **(A-N)** Levels of cytokines IL-6, IL-1β, TNF-α, IFN-γ, IL-17A, MIP-1α, MCP-3, GM-CSF, IP- 10, RANTES, MIG, IL-12p70, IL-18 and MCP-1 in BALF from mice challenged with LPS without 4-PBA treatment (*LPS*, n=14, graphed in red) and with 4-PBA treatment (*LPS + 4-PBA*, n=15, graphed in blue). Groups of unchallenged mice without 4-PBA (*C-*, n=6, graphed in black) and with 4-PBA treatment (*4-PBA*, n=9; graphed in green) were also evaluated. Colored lines and whiskers denote mean ± SEM for every data set. Hash marks indicate significant difference versus non-LPS challenge conditions (^#^*P* < 0.05, ^##^*P* < 0.01, ^###^*P* < 0.001) and a straight line between *LPS* and *LPS + 4-PBA* (\**P* < 0.05, \*\**P* < 0.01, \*\*\**P* < 0.001) by Two-way ANOVA followed by Tukey’s post-hoc test.

When animals challenged with LPS were treated with 4-PBA to reduce cellular stress we observed a significant rescue of several cytokine values. Among these, we observed a significant decrease in the three best documented general pro-inflammatory markers in COVID-19: IL-6, IL-1β and TNF-α (Figure 2A-C). These cytokines have been extensively related with bad prognosis in COVID-19 patients, being IL-6, as we aforementioned, one of the most important markers of deterioration of clinical profile and even associated with higher mortality rates^1, 58^. Other rescued cytokine values were detected in the macrophagic inducer IFN-γ (Figure 2D) and IL-17a, synthetized predominantly by CD4^+^ lymphocytes, strongly related with ARDS and responsible for neutrophil chemotaxis (Figure 2E). MIP-1α and MCP-3, produced initially by lung endothelial and epithelial cells at the beginning of the infection and by Mφ in later stages, also showed a reduction after treatment with 4-PBA (Figure 2F-G). Only one cytokine, GM-CSF, a myeloid growth factor associated with alveolar Mφ maturation, showed an increase after application of LPS and 4-PBA combined (Figure 2H). The remaining cytokines analyzed showed a slight decrease with LPS + 4-PBA compared to LPS alone without reaching statistical significance (Figure 2I-N).

In summary, the modulation of cellular stress with the use of 4-PBA showed the ability to influence the levels of cytokines related with monocytic/macrophagic activation and neutrophilia (IL-17a), suggesting a connection between cellular stress and certain immune lineages through the inflammatory response.

### The severity of the inflammatory response in ARDS is correlated with increased BiP in the alveolar space

As patient-derived data showed that increased BiP levels are correlated to risk factors and comorbidities of severe COVID-19, we next analyzed this stress marker in our ARDS mouse model. Results showed that LPS treatment increased BiP levels in the secretions of the alveolar space, and that this increase was ameliorated by 4-PBA treatment (Figure 3A). Furthermore, BiP levels had a significant and positive correlation with 12 of the 14 cytokines measured, including MCP-3, TNF-α, MIP-1α, IL-6 and IL-1β (Table 1), all the cytokines that were modulated by treatment with 4-PBA (Figure 2), further supporting that the hyper-inflammatory lung milieu is clearly connected with the ER stress response with participation of BiP.

**Figure 3.**
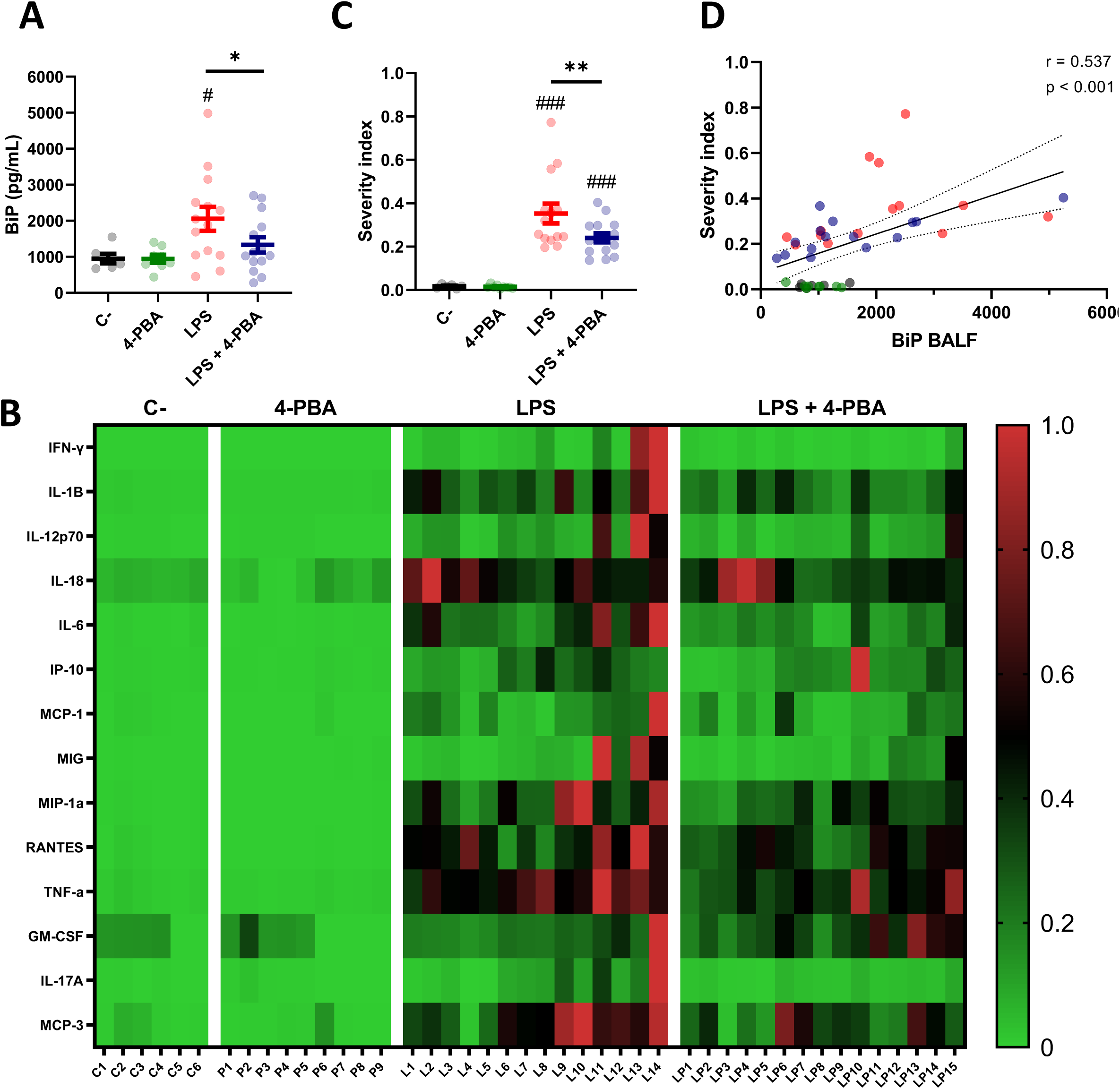
BiP levels correlate with ARDS severity. **(A)** BiP levels in BALF from control mice, 4-PBA treatment, challenged with LPS with and without 4-PBA treatment. Colored lines and whiskers denote mean ± SEM for every data set. Hash marks indicate significant difference versus non-LPS challenge conditions (^#^*P* < 0.05, ^##^*P* < 0.01, ^###^*P* < 0.001) and a straight line between *LPS* and *LPS + 4-PBA* (\**P* < 0.05, \*\**P* < 0.01, \*\*\**P* < 0.001) by Two-way ANOVA followed by Tukey’s post-hoc test. **(B)** Heatmap showing levels for all measured cytokines for every single mouse (In the X axis: C = *C-*; P = *4-PBA*; L = *LPS* and LP = *LPS + 4-PBA* with numbers indicating replicate number). Normalized cytokine values are depicted on a low-to-high scale (green-black-red). **(C)** Severity Index calculated as an average of normalized values for all cytokine by every single animal. Values near to 1 indicate more severe outcome whereas values tendent to zero a milder response. Colored lines and whiskers denote mean ± SEM for every data set. Hash marks indicate significant difference versus non-LPS challenge conditions (^#^*P* < 0.05, ^##^*P* < 0.01, ^###^*P* < 0.001) and a straight line between *LPS* and *LPS + 4-PBA* by Two-way ANOVA followed by Tukey’s post-hoc test. **(D)** Scatter plot showing a positive correlation between BiP levels in mice BALF and the calculated Severity Index tested by Pearson’s correlation coefficient.

To study how BiP could be linked with the severity of the ARDS, we analyzed cytokines levels from each animal individually to detect which mice suffered a stronger response to the LPS challenge and to determine responsiveness to the 4-PBA treatment. From this analysis, we created a Severity Index, calculated from the average value from the cytokines in each animal (Figure 3B-D). This severity index is, therefore, an indicative score of how strong the overall inflammatory response was by individual animals. Figure 3B shows how the majority of the cytokine highest values were found in LPS-treated mice, a group that included 13 of the 14 cytokine maximum levels in this experimental group. These qualitative observations were confirmed by the calculated Severity Index which was significantly higher in LPS challenged animals while significantly ameliorated by 4-PBA treatment (Figure 3C). Finally, we found a statistically significant correlation between increased levels of BiP in BALF with the Severity Index (Figure 3C and Table 1). Together, our data suggest a link between the severity of the inflammatory response and the ER stress state evidenced by increased BiP levels in BALF which can be modulated by the treatment with 4-PBA.

### Cell surface exposure of BiP is promoted in cell lineages responsible for the hyperinflammatory response

After we established that BiP is linked to inflammation and the severity of ARDS, we further studied the role of this chaperone in the immune cell environment responsible for the hyperinflammatory response. As previously mentioned, although BiP mostly resides in the ER, stress factors induce a translocation of BiP to the cell surface^59^. Furthermore, csBiP was shown to act as a coreceptor of several virus infections, including SARS-CoV-2^32, 33^. Therefore, we decided to evaluate the participation of pan-BiP or csBiP in ARDS. We first evaluated the mRNA expression and the protein levels of pan-BiP in lung tissues from our ARDS mouse model. Although we had detected an increase of available BiP in the alveolar space in response to LPS (Figure 3B), neither *Hspa5* gene expression (Figure S4A) nor whole protein abundance was significantly altered (Figure S4B-C), which suggests that the changes observed in BiP human serum and in the mice bronchoalveolar space are not correlated to changes in canonical ER stress.

Then, we looked into the cell surface BiP from lung tissue and since BiP-correlated cytokines pattern during inflammatory response in lungs is mainly orchestrated by neutrophils and monocytic lineages, we analyzed these cell populations from whole lung tissues (Figure 4A-L and Figure S5) and measured the levels of csBiP in all of them. At first glance, whole leukocyte population (CD45^+^ cells) did not change in number during the inflammatory process (Figure 4M Top), however, they showed a significant increase of csBiP (Figure 4M Bottom).

**Figure 4.**
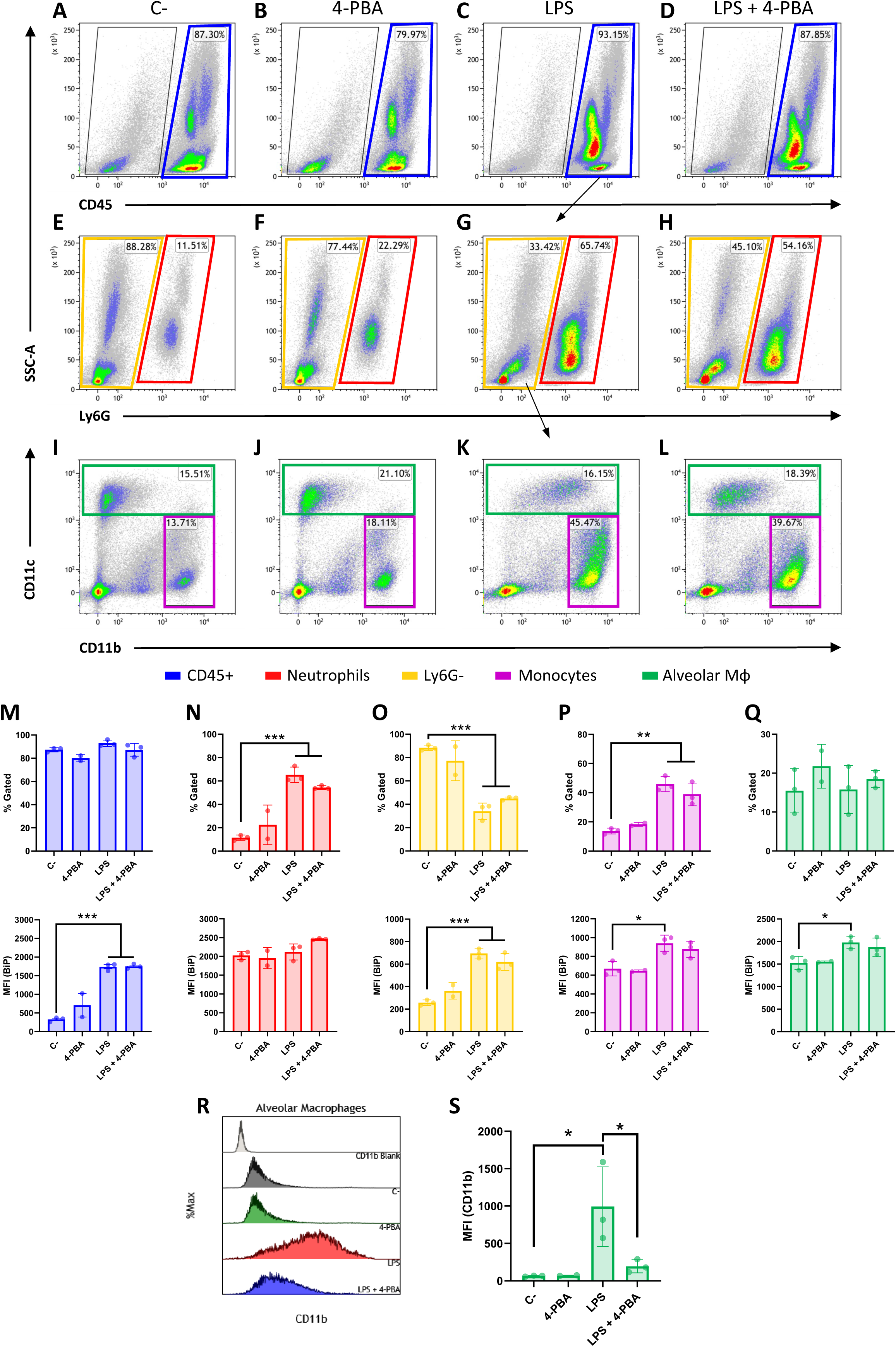
Cell surface BiP levels in immune lineages during the hyperinflammatory response. **(A-D)** Representative flow cytometry plots for CD45^+^ cells in blue squares. **(E-H)** Neutrophils are defined as CD45^+^ Ly6G^+^ in red squares. **(I-L)** Among the CD45^+^ Ly6G^-^ population in yellow squares, we defined alveolar macrophages and DCs (CD45^+^ Ly6G^-^ CD11c^+^) in green squares and monocytes as well as other myeloid phenotypes (CD45^+^ Ly6G^-^ CD11b^+^ CD11c^-/low^) in purple squares (n=3 mice per group, n=2 for “4-PBA” group). **(M-Q)** Percentage of gated cells and cell surface BiP levels measured by Median Fluorescence Intensity (MFI) of tagged csBiP antibody are represented in bar plots for every defined population. **(R-S)** Histogram graph show the intensity distribution of CD11b marker among Alveolar Mφ population **(R)** also represented as the average of its correspondent MFI in a bar plot **(S)**. All bar plots show mean ± SD for every treatment into the defined population. \**P* < 0.05, \*\**P* < 0.01, \*\*\**P* < 0.001 indicate statistically significant differences versus C-samples for a One-Way ANOVA with a Tukey’s multiple comparisons test.

When we analyzed each population independently, we found that neutrophil lung population (CD45^+^ Ly6G^+^) increased upon LPS stimulation, rising from 11.51% to 65.74% of the leukocyte population (Figure 4N Top). Interestingly, neutrophils showed the highest expression of csBiP amongst all the studied hematopoietic populations, although these levels of csBiP were not responsive to LPS stimulation (Figure 4N Bottom), indicating that they are naturally elevated in this cell population.

On the other hand, the non-neutrophilic population (Ly6G^-^ cells) showed lower basal levels of csBiP but a significant responsiveness to LPS stimulation, which increased 3 to 4 times compared to non-stimulated cells (Figure 4O). Within this non-neutrophilic population, we analyzed the monocyte subset identified as CD11b^+^ CD11c^-/low^, formed mainly by interstitial Mφ and residential monocytes^60^ (Figure 4I-L; purple square), whose population increased upon stimulation with LPS (Figure 4P Top). More importantly, this population showed increased levels of csBiP, when treated with LPS (Figure 4P Bottom). Finally, we analyzed the CD11c^+^ population formed by alveolar Mφ and dendritic cells (DCs). These cells did not increase in numbers with LPS treatment (Figure 4I-L, green squares, and Q Top) but similarly to monocytes, alveolar Mφ showed a significant increase in csBiP after LPS challenge (Figure 4Q Bottom).

Regarding LPS + 4-PBA treatments, even though we registered certain changes, there was no significant amelioration in the number of cells or csBiP translocation. Then, we wondered if 4-PBA modulated the immune activation state in any of these myeloid populations. For this, we analyzed CD11b expression levels, which is known to increase upon alveolar Mφ activation^61^ and we observed that this alveolar Mφ population was highly activated by LPS challenge while 4-PBA treatment rescued values of CD11b to normal levels (Figure 4R-S).

These results suggest that csBiP plays an important role in the modulation of the inflammatory response and that the two elevated immune cell populations increased in COVID-19 and ARSD, neutrophil and macrophages, are naturally elevated or have the ability to increase csBiP, further supporting their importance in the mediation of cellular stress during the hyperinflammatory response.

### A network of ER stress related proteins is altered during ARDS and crosstalk with pro-inflammatory factors

After establishing that BiP is involved in the ARDS mechanism of disease, we next analyzed the proteomic profile of lung tissue challenged with LPS and/or treated with 4-PBA to identify pathways and components that link BiP and cellular stress with the hyperinflammatory response. In LPS challenged lungs, we detected significant changes (p < 0.05 and Fold change > 1.5) in 159 proteins of the 3628 detected compared to negative control mice. String protein clustering identified four major clusters defined by GO term association (Figure S6A). Three of the four clusters identified were relatively expected: the first cluster contained proteins related to inflammation GO terms (37 proteins, Figure S6B); the second cluster included proteins related to interferon response (22 proteins, Figure S6C) and a third cluster included proteins from a more heterogeneous group related to cell metabolism and mitochondrial oxidative response (15 proteins, Figure S6D). More interestingly, the unsupervised algorithm also grouped a fourth cluster with 11 proteins classified under UPR and cellular stress GO-terms (Figure S6E). The existence of this differentially expressed cluster in the ARDS model suggests a solid participation of UPR-stress signaling in the mechanism of disease. Within this cluster, we did not find BiP, which showed no significant change in the proteomic analysis (Figure S7), consistent with our previous findings on the levels of pan-BiP in lung tissue (Figure S4). However, knowing that it is not pan-but csBiP the one involved in the modulation of ARDS and anti-stress treatment with 4-PBA, we studied interactions of proteins from this cluster with BiP (Hspa5) (Figure 5A-B). Among the proteins from the UPR/stress cluster, we found that BiP interacted with Hsph1, Hspa1a, Hspa1b, Bag3 (all molecular chaperones with a role in protein refolding and UPR signaling), Nup85 (a nucleoporin involved in CCR2-mediated chemotaxis of monocytes), Ripk1 (the Receptor-interacting serine/threonine-protein kinase 1, a key regulator of TNF-mediated apoptosis, necroptosis and inflammatory pathways, and the main connection between the UPR and inflammatory clusters; Figure S6A), S100a11 (a calcium binding protein inducible by ER stress) and H2-Q6 or HLA-G (a component of the Major Histocompatibility Complex I,G related to diseases like asthma, pre-eclampsia and to the antigen recognition of SARS-CoV-2^62^. These interactions revealed a network of proteins that connect a major stress pathway, the UPR signaling, to inflammation and infection.

**Figure 5.**
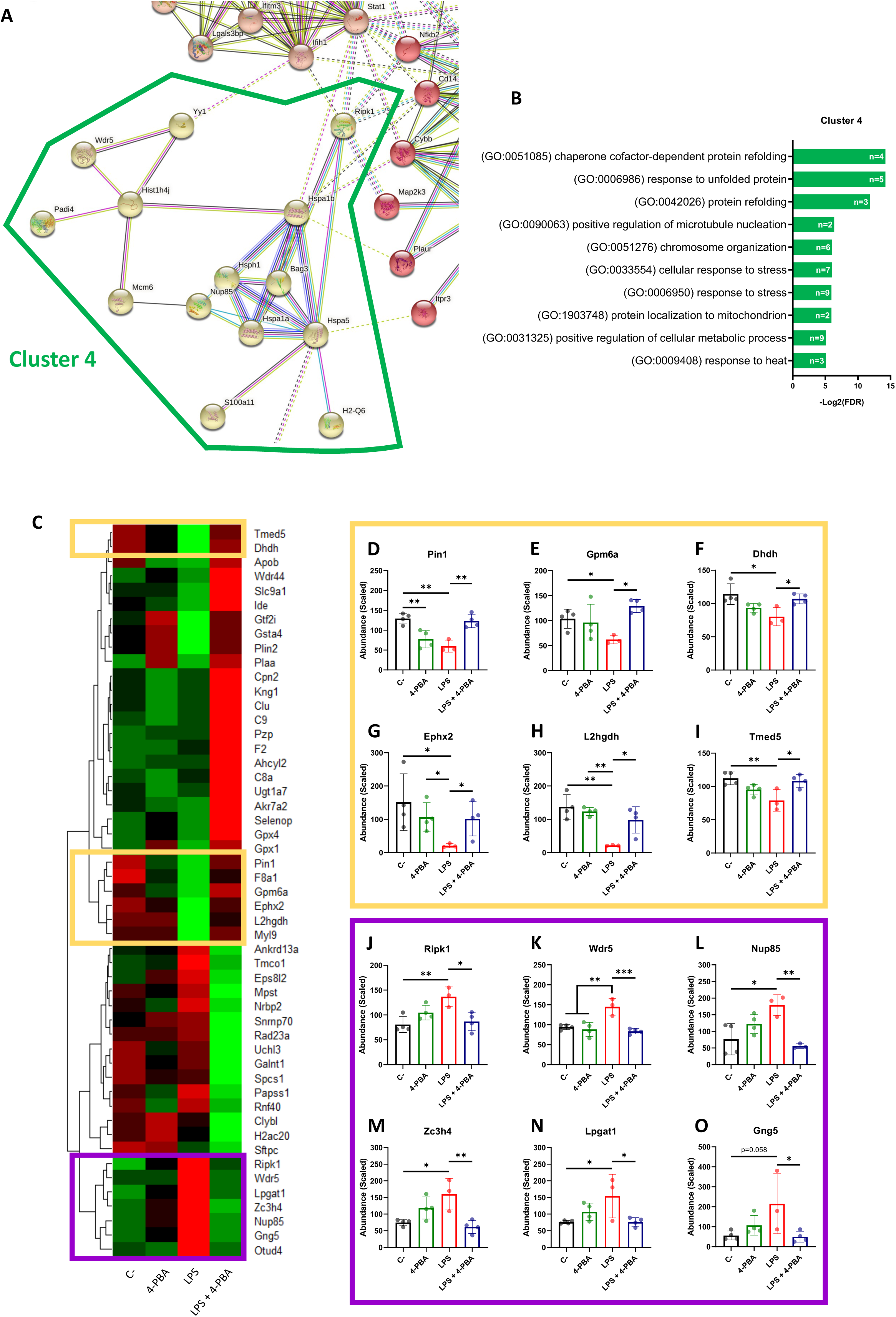
Differentially expressed proteins group as UPR/ER stress and inflammatory clusters linked by BiP and Ripk1. **(A)** StringDB network showing the associations between proteins differentially expressed in response to LPS challenge in mice lungs forming a cluster detected by an unsupervised Markov Cluster Algorithm (MCL). **(B)** Bar plots showing the Top-10 enriched Biological Processes associated with this cluster ordered by False Discovery Rate. Every single bar indicates the number of proteins associated with every GO-term. **(C)** Hierarchical clustered heatmap showing relative quantities of the 51 proteins expressed differentially in “*LPS*” group versus “*LPS + 4-PBA*” group. **(D-O)** Proteins with decreased (D-I, yellow squares) or increased (J-O, purple squares) levels after LPS challenge and that were rescued to normal levels after 4-PBA treatment. Bar plots show in detail the mean ± SD by treatment for every one of those highlighted proteins. \**P* < 0.05, \*\**P* < 0.01, \*\*\**P* < 0.001 indicate statistically significant differences between samples linked with a straight line for a One-Way ANOVA with a Tukey’s multiple comparisons test (n=4 for *C-*, *4-PBA* and *LPS + 4-PBA* groups; n=3 for *LPS* group).

As anti-stress treatments demonstrated to immunomodulate ARDS inflammatory response we next studied the proteomic changes between lungs challenged with LPS and LPS + 4-PBA to identify proteins involved in the amelioration of the inflammatory response. We found a group of 51 proteins with significant changes between LPS and LPS+4-PBA treated lungs (Figure 5C). We focused our attention in the ones whose levels changed upon LPS challenge and were then restored after 4-PBA treatment, and we identified 12 proteins that followed this pattern (Figure 5C-O). Levels of six of these proteins dropped with LPS and returned to normal values after 4-PBA treatment (Figure 5D-I): Pin1 (a peptidyl isomerase with a role in regulation of TP53, stress and cytokine signaling in immune system)^63^, Gpm6a, Ephx2, L2hgdh, Dhdh (general metabolic modulators) and Tmed5 (Transmembrane P24 Trafficking protein 5 involved in ER-Golgi trafficking and WNT signaling)^64^. The other six proteins had elevated levels upon LPS challenge and returned to low levels with 4-PBA (Fig. 5J-O): Ripk1 (the previously described key TNF regulator)^65, 66^, Wdr5 (WD Repeat Domain 5, a Cilia associated protein with GO-term related to histone modification and also present in cluster 4 in Figure S6E), Nup85 (the previously described nucleoporin involved in monocyte chemiotaxis)^67^, Zc3h4 (A Zinc Finger CCHtype Containing 4 protein involved in transcriptional regulation), Lpgat1 (the metabolic enzyme Lysophosphatidylglycerol Acyltransferase 1) and Gng5 (the G protein Subunit Gamma 5 related to immune response through CCR3 signaling)^68^. It is interesting to note that within this small group of proteins, three belong to the previously described UPR/Stress cluster (Ripk1, Wdr5, Nup85) and two to the inflammation cluster (Gng5, Lpgat1) (Figure 5), further suggesting the existence of a network of proteins that connect stress and inflammation. Particularly interesting is Ripk1, a kinase activated by SARS-CoV-2 infection in lungs that when inhibited reduces the viral load and mortality in COVID-19 humanized mouse model^69^. As observed in our proteomic studies, Ripk1 is the main link between ER stress and inflammation clusters and a direct interactor of BiP.

Overall, our work uncovers a connection between cellular stress represented by BiP and the hyper-inflammatory response induced in ARDS. It defines csBiP as a key modulator of immune lineages and as a biomarker of severity for respiratory infectious diseases such as COVID-19. We also establish a network of proteins that crosstalk between UPR/stress signaling and inflammation and demonstrate the potential of anti-stress therapies with chemical chaperones such as 4-PBA to treat ARDS related diseases.

## Discussion

Our research demonstrates a connection between inflammation and cellular stress through the UPR regulator BiP. Until recently, BiP has been defined as a chaperone assisting protein folding and UPR signaling within the ER compartment, however, this multifunctional protein has also been found to translocate to other cell locations expanding its role from an ER stress regulator to a general cellular stress transducer in the cytoplasm, mitochondria, and cell surface^28, 29^. Evidence that cell surface BiP influences ligand and antigen recognition is well documented^30, 70^, especially in COVID19, where BiP recognition by SARS-CoV-2 has been recently demonstrated^32, 71, 72^. This role, together with the fact that BiP reaches the cell surface upon stress stimulus, makes it a strong candidate to link inflammatory extracellular signals and stress in immune cells. This is supported by several studies that show that stress and inflammation pathways influence each other^21, 22, 45, 73–76^. Our results further support this connection by uncovering an inflammation-infection feedback system mediated by the ER stress regulator BiP. Initial infection-inflammatory process induces cellular stress, increasing cell surface BiP in immune lineages responsible for cytokine release and favoring virus entry through overexposed BiP, feeding the inflammatory process into a cytokine storm.

ER or cellular stress have also been related to the mechanism of disease of multiple pathologies^17, 23, 27, 73, 77, 78^, including clinical and subclinical manifestations classified as risk factors of COVID-19 (hypertension, diabetes, cardiovascular disease, obesity, autoimmune and respiratory diseases, among others). All these pathologies have been demonstrated to rise BiP levels and show signs of cellular stress, some of them even to be treatable by molecular chaperones^79, 80^. As a pre-existing state of cellular stress means abnormal levels of BiP, and BiP is able to feedback the inflammatory response^22, 75, 81^, it is no surprise that these risk factors push cells and tissues closer towards the molecular stress threshold that facilitates the hyperinflammatory response during ARDS.

Our results strongly support that BiP levels in blood or in bronchoalveolar fluid can be used as an early severity biomarker of risk of pneumonia in COVID-19 and other respiratory inflammatory diseases. Statistically, our study of almost 200 patients suggests that any SARS-CoV-2 positive patient that shows BiP values of 300pg/ml or higher in serum at the beginning of the infection has a 100% probability of developing pneumonia. This translates into a powerful prognosis tool, easy to apply in the clinic, however, it is important to note that it does not predict all patients that end up with a severe pneumonia, but a 19,48% (15 of the 77 patients with severe pneumonia in our cohort). Still, this 19,48% represents a risk group of patients where prognosis of development of severe pneumonia could have been applied with absolute certainty.

Experiments with the anti-stress agent 4-PBA indicate that stress does not act as a switch but as a modulator of the inflammatory response with a major significance in the transition from a moderate to severe respiratory disease. Most importantly, 4-PBA experiments suggest that ARDS and the hyperinflammatory response in lungs can be ameliorated by anti-stress drugs through small changes in the cytokine signaling pathways without blocking whole pathways that intervene in the immune response. What remains to be tested is its efficiency in avoiding the development of a severe pneumonia in patients with high levels of BiP in blood, but it is clear that 4-PBA is a strong candidate for the treatment of ARDS and that it seems like a viable option due to the fact that it is an approved drug.

There is still a certain gap in the knowledge about how the localization of BiP at the cell surface translates into changes in the cellular cascades that modulate cytokine pathways. From our proteomic studies, a potential candidate is RIPK1, an intermediary kinase between the UPR/stress proteins and the inflammation and the interferon response. Ripk1 interacts with both Hpsa5 (BiP) and Hpsa1a within this cluster, but it also interacts with Nfkb2 (Nuclear factor NF-kappa-B 2), which is present in many inflammatory and immune pathways^82, 83^. RIPK1 seems to act as a bridge between stress and the immune response also through its interactions with CD14 (Monocyte differentiation antigen CD14 (Cd14) that mediates the immune response to bacterial LPS), MAP2K3 (which is activated by cytokines and environmental stress processes), STAT1 (Signal transducer and activator of transcription 1 (Stat1) which modulates responses to many cytokines and interferons^84^) and IFIH1 (Interferon-induced helicase C domain-containing protein 1 (Ifih1) that acts as a viral sensor and plays a major role in the activation of the antiviral response through an increase in pro-inflammatory cytokines and type I interferons^85^) from clusters 1 and 2 (Figure 5 and S6). This points RIPK1 as a strong candidate to mediate BiP signal transduction in the cell surface of neutrophils and alveolar macrophages while acting as a co-receptor of NFKB/TNF signals^86, 87^. Recently, RIPK1 activation was described in human COVID-19 lung samples; inhibition of RIPK1 with the use of small molecules reduced lung viral load and mortality in ACE2 transgenic mice^69, 88^. This further supports that the reduction of BiP levels could result beneficial for the treatment of patients with severe COVID-19. Another interesting interactor of BiP during inflammation of ARDS is H2-Q6, a histocompatibility factor that has been related to SARS-CoV-2 recognition^62, 89^. This suggests that BiP and H2-Q6 could be favoring virus recognition.

In summary, our research connects stress and inflammation during ARDS in diseases such as COVID-19, it finds a valuable early biomarker of severe pneumonia, suggests a mechanism of severity by csBiP exposure in immune lineages and offers proof-of-concept for a new therapeutic approach through the use of anti-stress drugs.

## Methods

### Patients

Human serum was obtained from 194 confirmed positive patients for SARS-CoV-2 by clinical qPCR test (108 male and 86 females with a mean age of 64.85 ± 16.25 years; ranging between 0 and 94 years). The whole blood samples were collected at the time just after hospital admission at the beginning of the SARS-CoV-2 infection. This cohort included patients with different degree of clinical severity, from asymptomatic to lethal COVID-19 from the first wave of the pandemic (March-June 2020).

We also enrolled 30 healthy patients without any known comorbidities in order to determine the normal range of BiP in blood to compare with COVID-19 patients.

More detailed information about blood donor selection/exclusion criteria can be found in supplementary document 1.

### LPS challenge and 4-PBA treatment

Male wild-type C57BL/6J mice between 8-12 weeks old were used in this study (Charles River laboratories). Mice were kept on a 12h light/dark cycle with free access to food and water. All procedures and animal care follow the guide for the care and use of laboratory and all experimental protocols were approved by safety and ethics committees of the IBIMA-Bionand Platform Institute for the Animal Research Facility.

Before the challenge, mice were anesthetized with a mixture of ketamine and xylazine (50 and 4 mg/Kg, respectively). Then, 2 mg/Kg of Lipopolysaccharides (LPS) from *E. coli* O111:B4 (L2630, Sigma) diluted in PBS were intranasally administered in 50 µL of total volume (≈ 50 µg by animal). Procedure for control mice was identical but instillation was performed with PBS. Then, 200 mg/Kg of Sodium 4-phenylbutirate (4-PBA) (kindly provided by Scandinavian Formulas) diluted in PBS were administered intraperitoneally at 0, 6 and 8 hours after LPS challenge. Animals that did not receive 4-PBA were treated with the corresponding PBS volume in the same way.

Mice samples were harvested 24 hours after LPS instillation. Mice were euthanized by inhaled isoflurane overdose. Abdominal cavity was opened, blood was collected from the abdominal vena cava and transferred to EDTA pretreated tubes (BD Vacutainer®). Then, the trachea was exposed to perform a bronchoalveolar lavage (BAL) with 800 µL of chilled PBS which was carefully infused into the lungs and withdrawn three times. The collected fluid was then centrifuged at 800 x*g* during 10 minutes at 4 °C. Supernatant (BALF) was stored at -20 °C until posterior measurements. After that, washed lungs were extracted, snap-freeze in liquid nitrogen and stored at -80 °C for posterior molecular and proteomic analyses. Hematological analyses were immediately performed in a DF50 DYMIND hematology analyzer following the manufacturer instructions.

### Cytokines content in BALF

Cytokines content in mice BALF were measured by a custom designed ProcartaPlex multiplex immunoassay (Invitrogen) including the mice analytes: IL-1β, TNF-α, IL-6, IFN-γ, IL-17A, MIP-1α, MCP-3, GM-CSF, IP-10, RANTES, MIG, IL-12p70, IL-18 and MCP-1. Assays were performed following manufacturer instructions using undiluted BALF samples. Measurements were done in the Bio-Plex 200 system and calculations of cytokine content were performed in Bio-Plex Manager 6.0 software (Bio-Rad).

### ELISA

Human circulating BiP from serum samples and BiP content in mice BALF samples were evaluated with commercially available ELISA kits (LS-F11578 and LS-F17959 respectively, from LSBio). Human samples were diluted 1:5 in the supplied buffer whereas mice BALF were performed undiluted. Every single sample and standard were measured in duplicate.

### RT-qPCR

RNA was extracted from lung tissue previously broken up with a mortar and pestle using TRIzol^TM^ Reagent (catalog no. 15596026, Thermofisher). Complementary DNA (cDNA) was prepared from 1 μg of RNA using PrimeScript^TM^ RT Master Mix (catalog no. RR036A, Takara). qPCR was performed using TB Green Premix Ex Taq^TM^ (catalog no. RR420L, Takara). Gene expression was calculated using the 2ΔΔCT method of analysis against the stable housekeeping gene TBP. Five biological replicates were performed with three technical replicates each. qPCR primers were: BiP, 5’-TGAAACTGTGGGAGGAGTCA-3’ (forward), 5’-TTCAGCTGTCACTCGGAGAA-3’ (reverse), TBP, 5’-AGAACAATCCAGACTAGCAGCA-3’ (forward), 5’-GGGAACTTCACATCACAGCTC-3’ (reverse).

### Western blot

For protein analyses, lung tissue was mechanically broken up with a mortar and pestle and collected in IP lysis buffer (catalog no. 87787, Thermo Fisher Scientific) supplemented with proteinase inhibitors. Concentrations were determined using the Pierce^TM^ BCA Protein Assay Kit (catalog no. 23227, Thermofisher).

For Western blot analyses, protein lysates were separated by electrophoresis on 10% SDS–polyacrylamide gels, transferred to polyvinylidene difluoride membranes, blocked in 5% milk, and probed with primary antibodies: anti-BiP antibody (1:1000; catalog no. 3177, CST), anti-GAPDH (1:2000; catalog no. 2118). Peroxidase-conjugated secondary antibodies (catalog nos. 7071 and 7072, CST) were used, and immunocomplexes were identified using the ECL (enhanced chemiluminescence) Detection Reagent (catalog no. 322009, Thermofisher). Fiji was used to quantify bands after gel analysis recommendations from ImageJ and (http://rsb.info.nih.gov/ij/docs/menus/analyze.html#gels).

### Proteomic analysis by label-free quantification-based mass spectrometry

Mice lungs (n=4 for every treatment group) were mechanically broken up with a mortar and pestle and sonicated for 30 minutes in RIPA buffer to obtain protein extracts. After quantification by BCA method, volumes were adjusted to equalize all concentrations (One sample of the group “LPS” was excluded at this level by abnormally low values). The carried-out protocol was previously described in detail^90^ and adapted for lung tissue. Briefly, proteins were stacked in an acrylamide gel, bands were cut and further treated to be reduced with DTT, carbamidomethylated and digested with trypsin overnight. Then, resulting peptides were extracted, purified and concentrated for next steps and posterior mass spectrometric analysis.

Peptides samples were separated by liquid chromatography in Easy nLC 1200 UHPLC system coupled to a spectrometer coupled to a hybrid quadrupole-linear trap-Orbitrap Q-Exactive HF-X mass spectrometer for the analysis. Protein identification was performed, against the *Mus musculus* protein database of the SwissProt. Raw acquired data were analyzed on the Proteome Discoverer 2.4 platform (all by Thermo Fisher Scientific). Label-free quantification was implemented using the Minora function, setting the following parameters: maximum alignment retention time of 10 min with a minimum signal/noise of 5 for feature linkage mapping. The calculation of the abundances was based on the intensities of the precursor ions. Protein abundance ratios were calculated directly from the pooled abundances. p-Values were calculated by ANOVA based on the abundances of individual proteins or peptides. Only proteins with changes higher than 1.5-fold and with p-Values < 0.05 were considered significantly affected by the different treatments.

Protein-protein interaction of those significantly changed by LPS treatment against the control samples were analyzed and visualized using the online STRING database in order to establish a representative protein network associated with the response to endotoxin insult. Clustering was performed in the same website using the unsupervised MCL clustering tool with an inflation parameter = 1.4^91^. The resultant GO-terms list for the enriched biological processes in every single cluster were ascendent ordered by false discovery rate (FDR) and processed in the Revigo website tool in order to summarize it and removing redundant GO terms^92^.

### Flow cytometry

For flow cytometry, independent groups of mice were treated and euthanized following the same aforementioned procedure. In this case, they were bled cutting the inferior vena cava and both lungs were dissected. In these lungs the BAL was not performed in order to maintain the whole interstitial and alveolar populations to be processed for flow cytometry.

Harvested tissues were minced by scissors and digested in DMEM Low glucose (Sigma) + collagenase A (1 mg/mL) (Sigma) + DNase (0.05 mg/mL) (Roche) for 30 minutes at 37 °C in constant orbital agitation. Then, samples were vortexed for 10 seconds and passed through a 70 µm cell strainer to be disaggregated and erythrocytes were removed by incubation with ACK lysis Buffer (Sigma).

After extensive washing in Cell Staining Buffer (BioLegend), obtained single cell suspensions were stained with the fixable viability Zombie Aqua™ dye according with the manufacturer instructions (#423101; BioLegend, 1:500).

Then, all samples were treated with anti-CD16/32 (#14-0161-82; 0.5 µg/test, 10 min at 4 °C) for Fc-receptor blockage prior to staining procedure. Cells were incubated at 4 °C in the dark for 20 minutes with the following antibodies: Alexa Fluor 488 conjugated anti-GRP78 (#PA1-014A-A488, 1:50), eFluor™ 450 conjugated anti-CD45 (#48-0451-80, 1:100), Alexa Fluor 700 conjugated anti-CD11c (#56-0114-80, 1:50), Allophycocyanin (APC) conjugated Ly6G (#17-9668-80, 1:200) and phycoerythrin (PE) conjugated CD11b (#12-0112-81, 1:100). All the antibodies in this section were purchased from Thermo Fisher Scientific.

Finally, cells were fixed with 4% fresh formaldehyde at RT for 15 minutes, washed extensively and resuspended in Cell Staining Buffer to be evaluated on a BD FACS Aria Fusion flow cytometer (BD Biosciences). Results were analyzed with the software Kaluza (Beckman Coulter). Single stained and FMOs controls were included for every single antibody and for the viability marker in order to make the correspondent compensations and to determine all the cell population gates. Gating strategies are shown in supplementary figure S5.

### Statistical analysis

All statistical analyses were performed using GraphPad Prism and SigmaPlot 11.0. Unless stated different, all data are presented as mean ± SEM. Two tailed Student’s t-test, two-way ANOVA or ordinary one-way ANOVA with Tukey multiple-comparison test were performed for statistically significant differences among samples. Scatter plot were analyzed by Pearson’s correlation coefficient (r and its related *P*-value). Bold line shows the linear regression between the two variables and dotted lines denote the 95% confidence interval. Data sets from mice experiments of cytokines and BiP measurements were evaluated with the ROUT method (Q = 1%) to identify and exclude outlier values from nonlinear regressions^93^.

For proteomic analyses, results for *Label Free* protein quantification were generated by the Central Research Support Services from the University of Malaga. From that data, *abundance ratio* between samples from every treatment (calculated as a pairwise ratio) and their associated *p-Values* (from ANOVA *Background-Based* method) were used to classify proteins that significantly change in response to different treatments (Fold change > 1.5 and p-Value < 0.05). Volcano plots, hierarchical clustered heatmaps and correlation plots were performed in RStudio software.

## Acknowledgments

We thank patients from our cohort for their disposition to participate in this study and clinicians that diligently noted all the parameters included in this study. We would also like to thank the Scandinavian Formulas Inc. for kindly providing 4-PBA. Authors thank Casimiro Cardenas (Proteomics service at UMA) and David Navas (Flow cytometry service at UMA) for their technical assistance. This study was supported by Junta de Andalucía (Consejería de Salud y Familia) through CV20-81404 and PIGE-0178-2020. Universidad de Málaga and IBIMA-Plataforma Bionand funds (Plan propio UMA and IBIMA-TECH); Ministerio de Ciencia, Innovación y Tecnología (PID2020-117255-RB100). FC is supported by PCI2021-122094-2B and JMPT by FPU19/06951.

## Author contributions

Conceptualization: FC, JB and ID; Investigation: GR, OPP, SE; Writing: GR, FC and ID; Funding Acquisition: FC, JB and ID; Clinical data acquisition: OPP, LV and JLR; Mouse model optimization and data acquisition: DV, MC, DBV, and JMPT; Supervision: FC and ID.

---

**Supplementary Figure 1.**
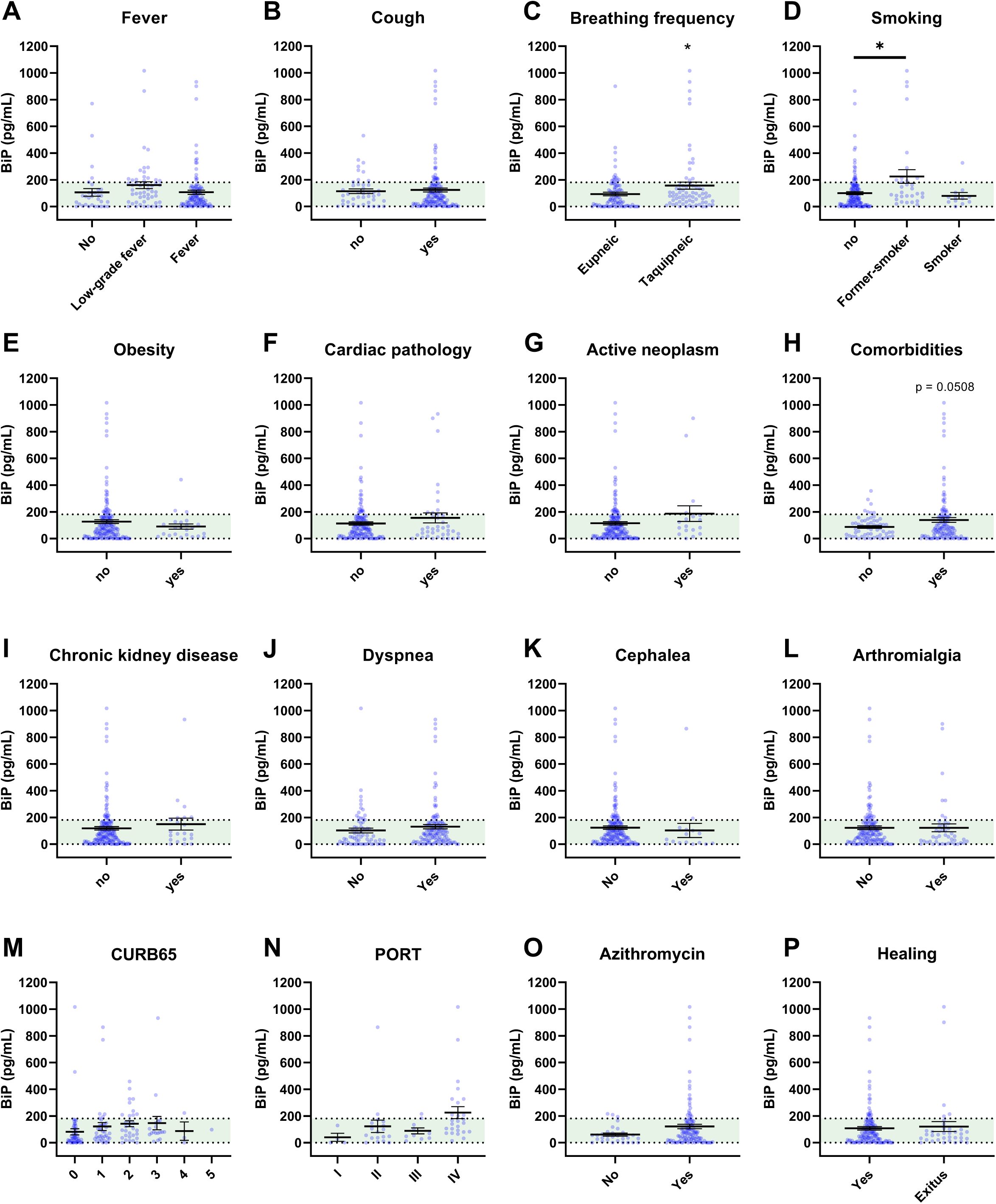
**(A-P)** Serum BiP levels classified by group of patients/donors. Black lines and whiskers denote the mean ± SEM of every data set. Green areas were defined between 5^th^ and 95^th^ percentiles of healthy donor’s data set as normal BiP levels in serum (0 and 181 pg/mL, respectively). \**P* < 0.05, \*\**P* < 0.01, \*\*\**P* < 0.001 indicate statistical significant differences between samples for a Two-Tailed unpaired t-Test **(A-C, E-L, O,P)** and One-Way ANOVA with a Tukey’s multiple comparisons test **(D, M, N).**

**Supplementary Figure 2.**
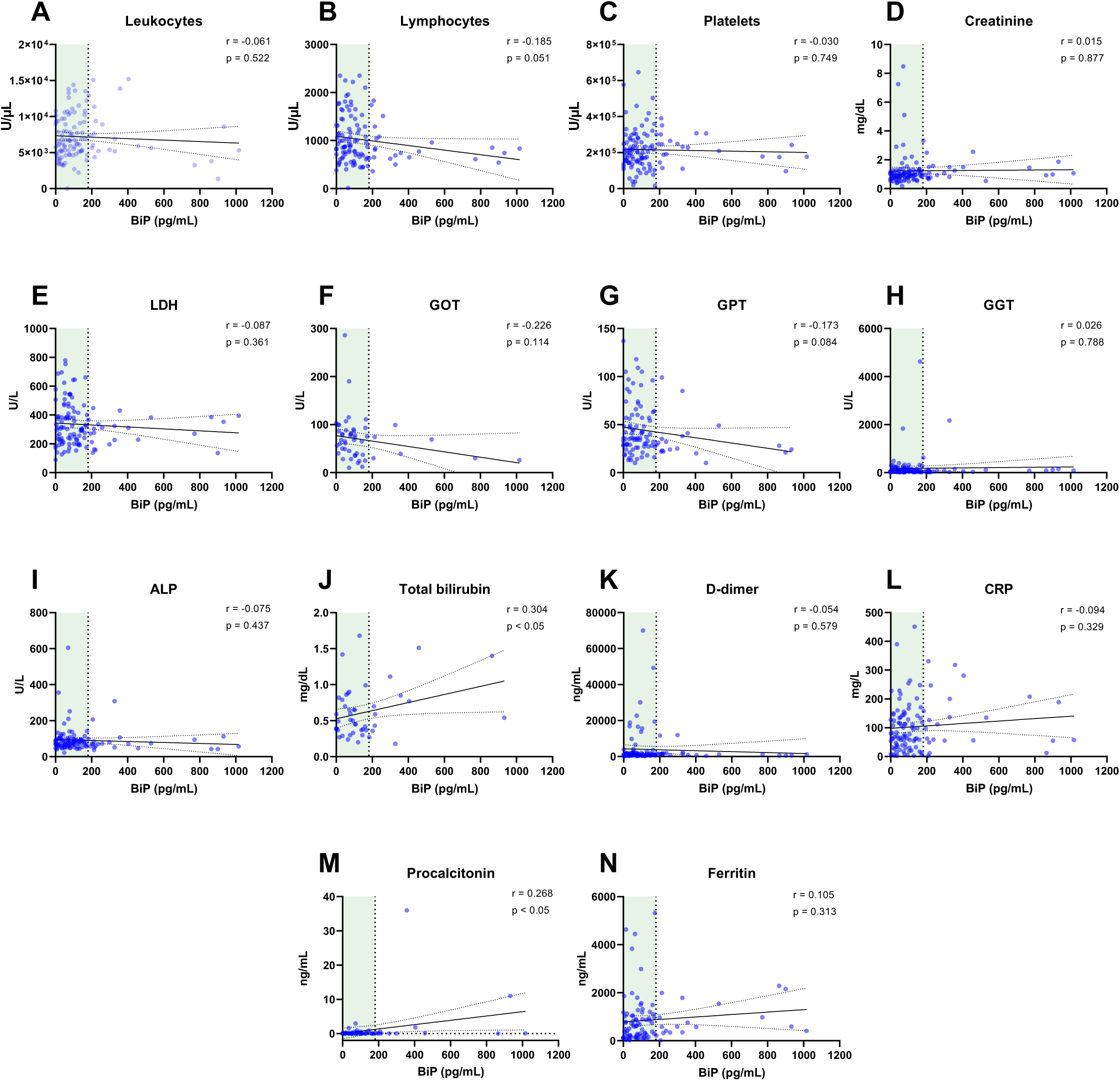
**(A-N)** Scatter plots showing the correlation between BiP levels versus different hematological and biochemical parameter levels in COVID-19 patient’s blood serum tested by Pearson’s correlation coefficient (r and its related *P*-value). Bold line shows the linear regression between the two variables and dotted lines denote the 95% confidence interval.

**Supplementary Figure 3.**
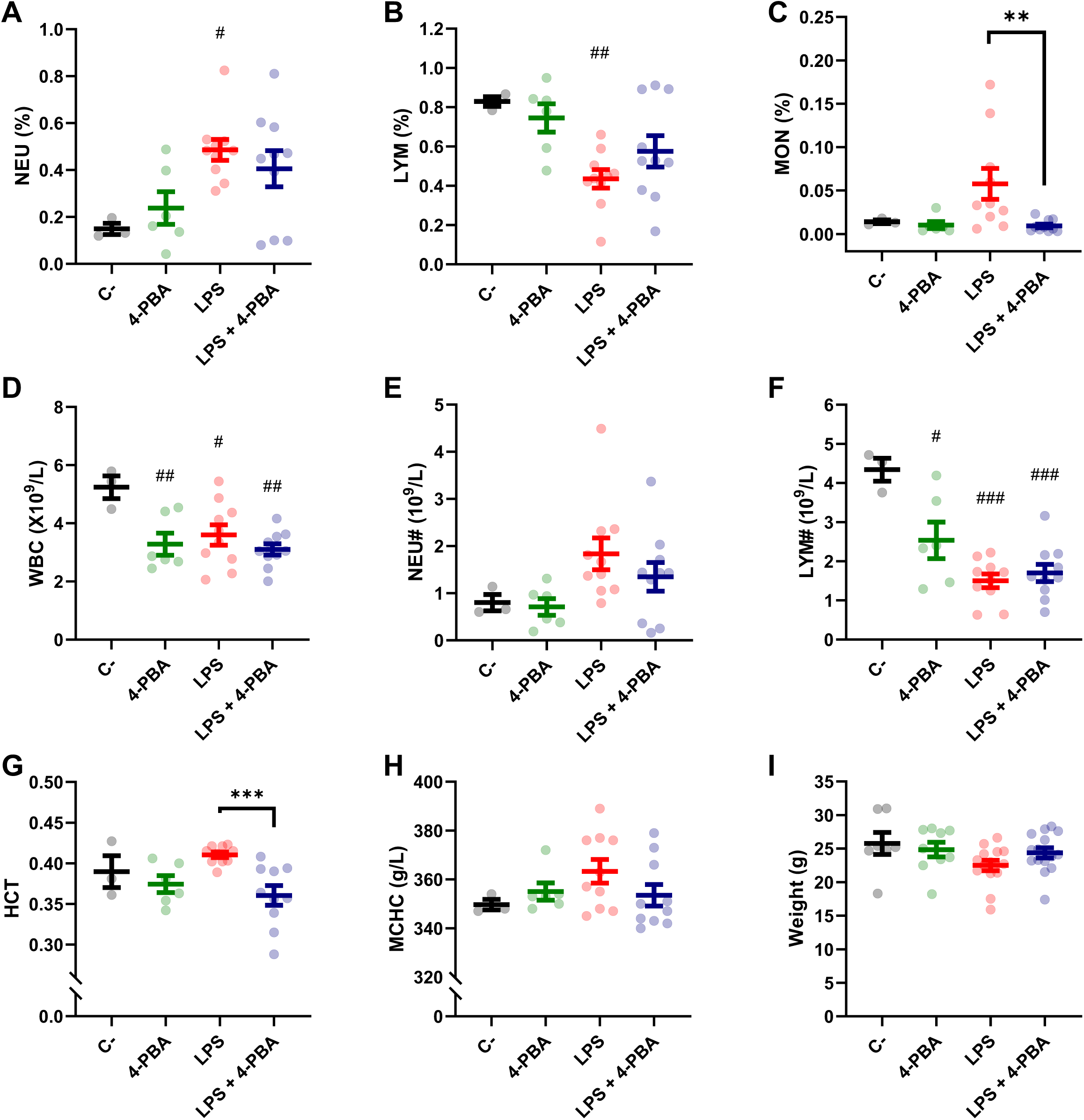
**(A-H)** Hematological analyses: percentages of neutrophils **(A)**, lymphocytes **(B)** and monocytes **(C),** total numbers of white blood cells (WBC) **(D),** total numbers of neutrophils **(E)** and total numbers of lymphocytes **(F)**, hematocrit (HCT) **(G)** and mean corpuscular hemoglobin concentration (MCHC) **(H)** in total blood obtained from mice challenged with LPS without 4-PBA treatment (*LPS*, n=10, graphed in red) and with 4-PBA treatment (*LPS + 4-PBA*, n=10, graphed in blue). Groups of unchallenged mice without 4-PBA (*C-*, n=3, graphed in black) and with 4-PBA treatment (*4-PBA*, n=6; graphed in green) were also evaluated. **(I)** Animal’s weight for every experimental group. Colored lines and whiskers denote mean ± SEM for every data set. Hash marks indicate significant difference versus C- (^#^*P* < 0.05, ^##^*P* < 0.01, ^###^*P* < 0.001) and asterisks between samples linked by a line (\**P* < 0.05, \*\**P* < 0.01, \*\*\**P* < 0.001) for a Two-way ANOVA followed by Tukey’s post-hoc test.

**Supplementary Figure 4:**
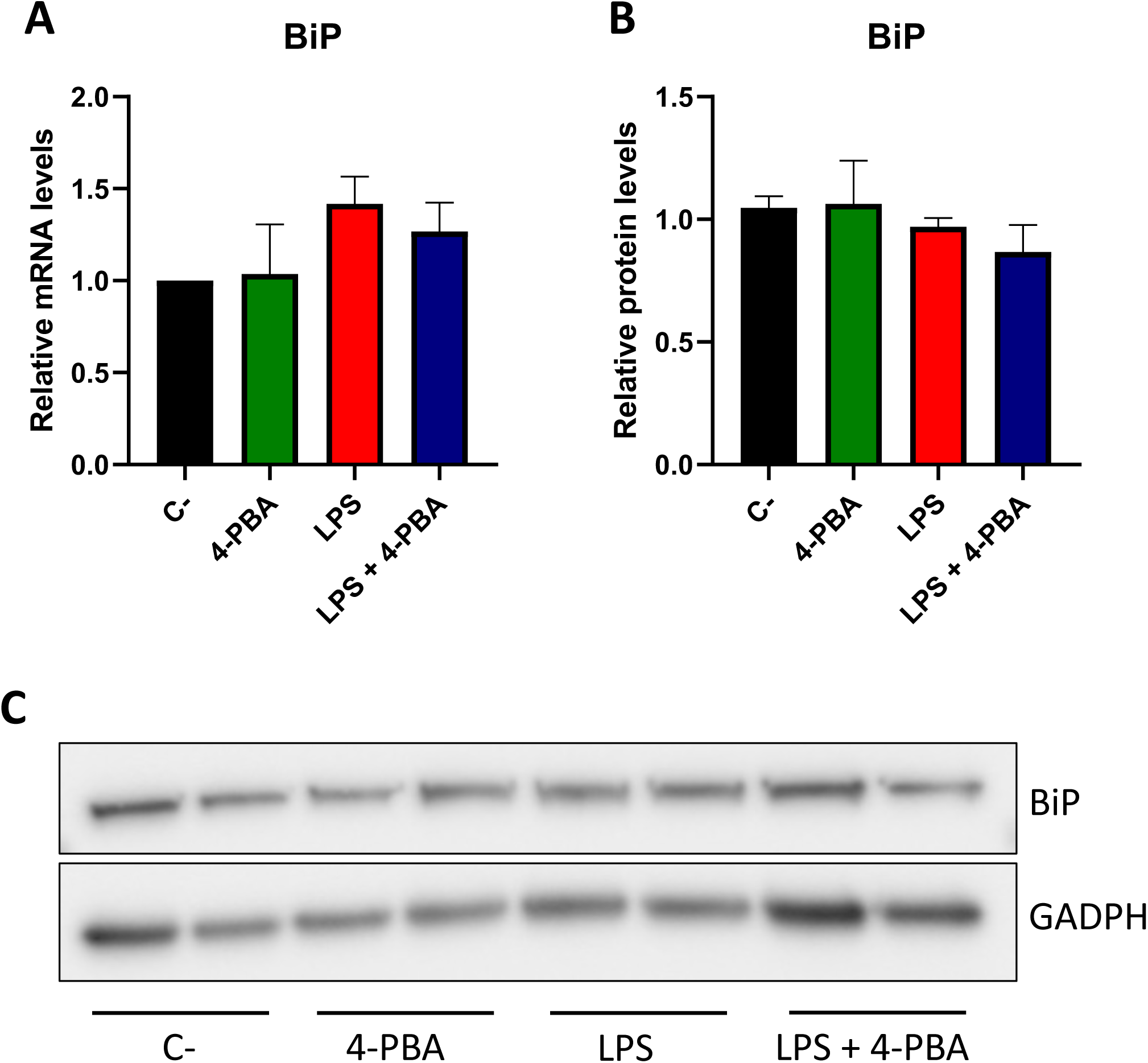
**(A)** Quantitative gene expression of Hspa5 (BIP) measured by RT-qPCR. **(B-C)** Total (pan) levels of BiP protein in lung tissues; western blot **(B)** and corresponding quantification of protein levels representation **(C).** No statistical significance was found among the represented conditions.

**Supplementary Figure 5.**
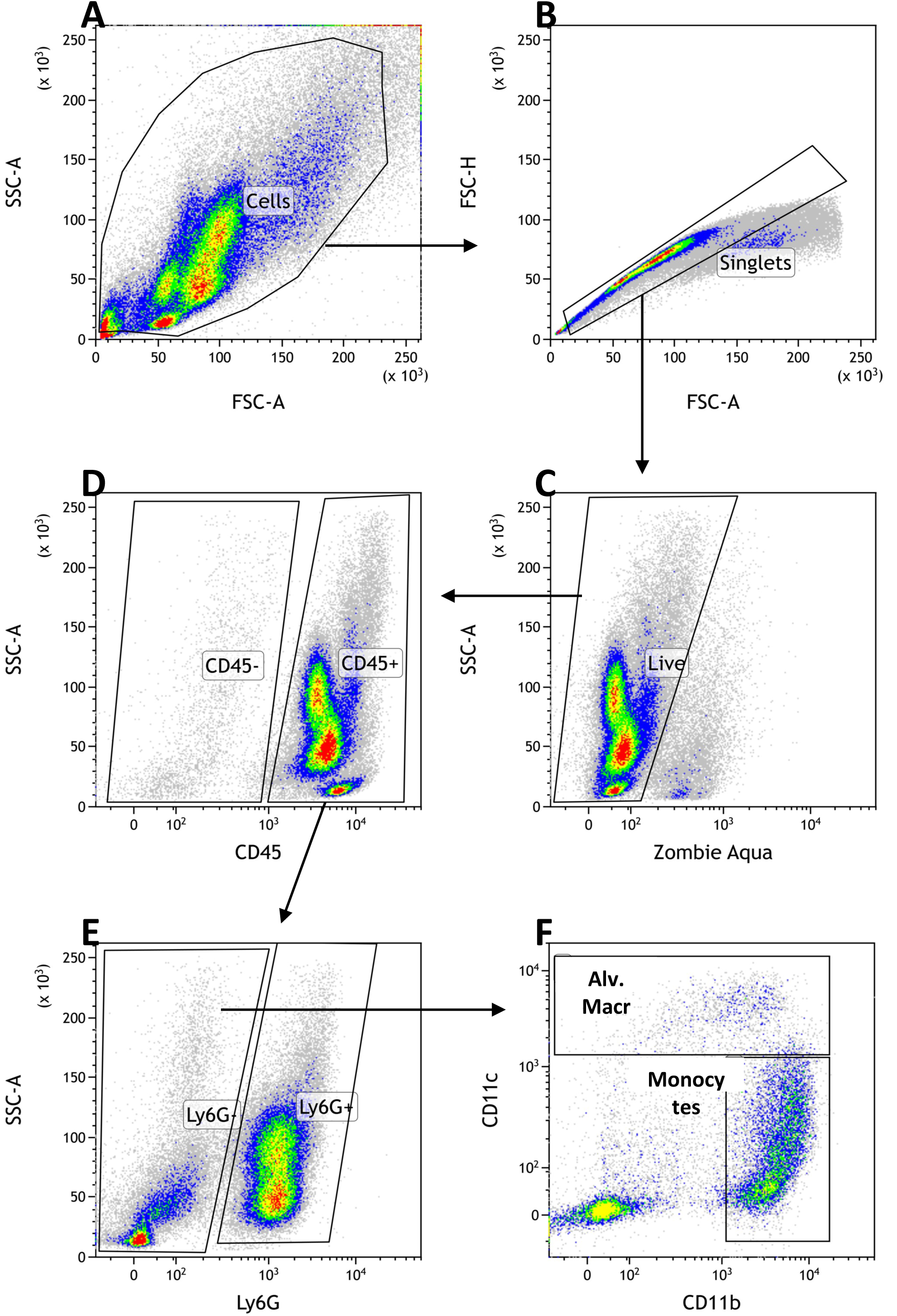
Flow cytometry gating strategy. **(A)** Representative scatter plots showing FSC-A x SSC-A gating to exclude debris based on size and granularity; **(B)** FSC-A x FSC-H to exclude doublets. **(C)** Zombie Aqua^Tm^ fixable viability marker to identify live cells (negative for the marker. **(D)** Staining with CD45 to identify hematopoietic cell linages. **(E)** Ly6G was used to identify neutrophils (as Ly6G^+^). **(F)** Among the Ly6G^-^ cells, CD11b x CD11c was used to identify alveolar macrophages and dendritic cells population (as CD11c^+^ with variable levels of CD11b) and monocytes as long as other myeloid subsets (as CD11b^+^ CD11c^-/low^). Gates for all viability marker and all antibodies were determined using respective fluorescence minus one (FMO) control.

**Supplementary Figure 6.**
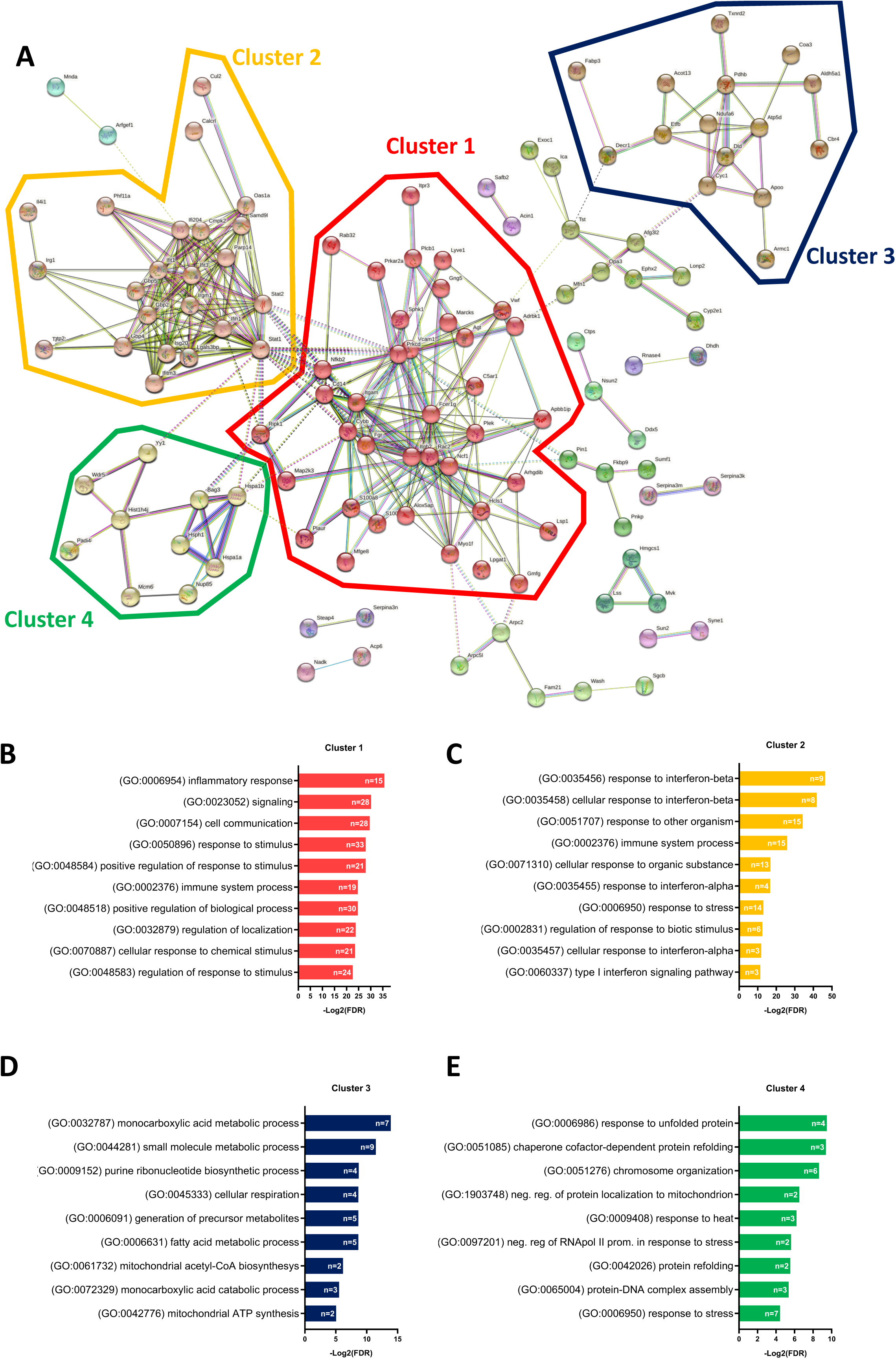
**(A)** StringDB network (without inclusion of Hspa5) showing the associations between proteins differentially expressed in response to LPS challenge in mice lungs forming 4 principal clusters detected by an unsupervised Markov Cluster Algorithm (MCL). **(B-E)** Bar plots showing the Top-10 enriched Biological Processes associated with every cluster ordered by False Discovery Rate. Single bars indicate the number of proteins associated with every GOterm.

**Supplementary Figure 7.**
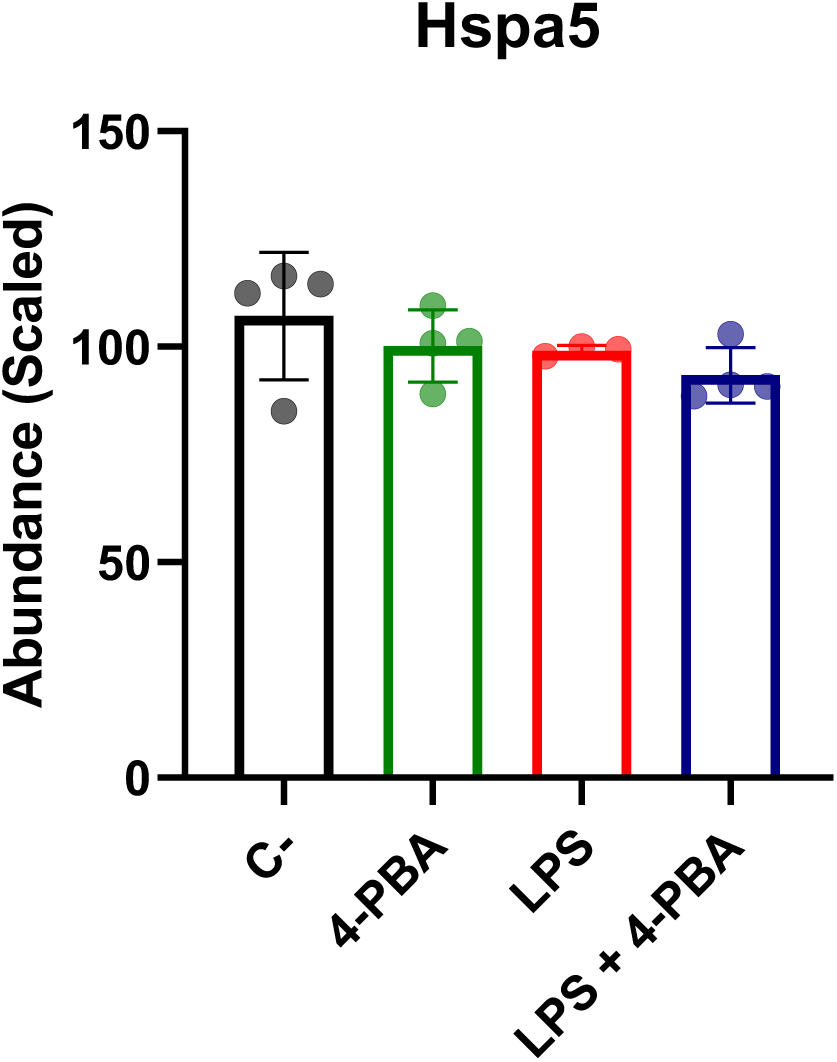
BiP Levels measured by proteomic analysis in lung tissues showing the mean ± SD of the scaled abundances. There were no statistically significant differences between samples for a One-Way ANOVA with a Tukey’s multiple comparisons test (n=4 for *C-*, *4-PBA* and *LPS + 4-PBA* groups; n=3 for *LPS* group).

**Supplementary Table 1:** Pneumonia severity categories

| Severity | Oxygen requirement | Parametric values |
| --- | --- | --- |
| No pneumonia | No | - Saturation >94%<br>- Eupneic * |
| Mild severity | Low Flow | - Saturation <94%<br>- Eupneic * |
| Moderate severity | Low Flow | - Saturation 90%-94%<br>- Possible Tachypnea**<br>- Risk factors identified |
| High severity | High Flow<br><u>Noninvasive mechanic ventilation</u> | - Saturation <90%<br>- Possible Tachypnea**<br>- Risk factors identified |
| Very high severity | <u>Invasive mechanic ventilation</u> | - Saturation <90%<br>- <u>Tachypnea</u><br>- Risk factors identified |

\* Eupneic <22RPM
\*\* Tachypnea (22 O > 22 RPM)

